# Sweet and sticky: increased cell adhesion through click-mediated functionalization of regenerative liver progenitor cells

**DOI:** 10.1101/2024.06.21.599861

**Authors:** Amaziah R. Alipio, Melissa R. Vieira, Tamara Haefeli, Lisa Hoelting, Olivier Frey, Alicia J. El Haj, Maria C. Arno

## Abstract

The burgeoning field of cell therapies is rapidly expanding, offering the promise to tackle complex and unsolved healthcare problems. One prominent example is represented by CAR T-cells, which have been introduced into the clinic for treating a variety of cancers. Promising cell-based candidates have also been developed to promote tissue regeneration, showing high potencies for the treatment of damaged liver. Nevertheless, in the remit of regenerative medicine, cell-therapy efficacies remain suboptimal as a consequence of the low engraftment of injected cells to the existing surrounding tissue. Herein, we present a facile approach to enhance the adhesion and engraftment of therapeutic hepatic progenitor cells (HPCs) through specific and homogeneous cell surface modification with exogenous polysaccharides, without requiring genetic modification. Coated HPCs exhibited significantly increased markers of adhesion and cell spreading, demonstrating preferential interactions with certain extra-cellular matrix proteins. Moreover, they displayed enhanced binding to endothelial cells and 3D liver microtissues. This translatable methodology shows promise for improving therapeutic cell engraftment, offering a potential alternative to liver transplantation in end-stage liver disease.

---

Liver transplantation currently stands as the only therapeutic option for multiple acquired and congenital liver diseases.^1–4^ However, the paucity of liver donors limits this option to a minority of patients, highlighting a need for alternative treatment strategies.^5,6^ Cell-based regenerative therapies may circumvent the requirement for organ donation and invasive organ transplantation through the *ex vivo* expansion and reintroduction of regenerative cells.^7,8^ Pre-clinical *in vivo* models have consistently demonstrated remarkable levels of functional restoration at sites of engraftment within failing livers.^9–11^ Despite these promising results, the challenge of low engraftment remains a significant hurdle, impeding the broader adoption of cell-based therapies in clinical practice.^8^ Specifically, nearly 80-90% of transplanted hepatocytes are destroyed, also as a consequence of inadequate adhesion to the sinusoidal endothelium,^12^ with hepatic progenitor cells (HPCs) engrafting even less (< 5%).^13^ Thus, there is a clear need for exploring novel methods to increase cell engraftment and improve the overall efficacy of cell-based therapies.

Cellular engraftment is mediated by the cell membrane and the immediate pericellular matrix, making the cell surface a rational target for modification. Genetic modification (GM) strategies that alter the expression of surface receptors represents a traditional approach for site-specific homing of regenerative cells.^14,15^ However, these are often time-consuming and costly, carrying substantial risks of insertional mutagenesis and adverse heterogenous expression.^16–18^ As a result, alternative strategies have been developed to expand the plethora of foreign functionalities that can be conjugated to the cell surface, including bioactive and therapeutic payloads that can modify cell behavior and cell interactions with the surrounding environment.^19–24^ Among these, metabolic oligosaccharide engineering (MOE) pioneered by Bertozzi and co-workers has emerged as a promising technique for the integration of abiotic chemical functionalities onto cell surface glycans, which provide an ideal binding site for the attachment of macromolecules due to their high abundance at the cell surface.^25–29^

MOE employs unnatural sugar analogues bearing bio-orthogonal chemical handles to ‘hijack’ the cytosolic enzymes of the glycan biosynthetic pathway, effectively replacing naturally occurring sugars. The highest cell surface coverage can be achieved using *N*-acetyl mannosamine (ManNAc) derivatives, such as *N*-azidoacetylmannosamine-tetraacetate (Ac_4_ManAz), owing to their participation in the sialic acid biosynthetic pathway and efficient conversion to azidoacetyl sialic acids, which are incorporated into *N*- and *O*-linked sialoglycoconjugates.^28–32^ Using this technology, the membrane of mammalian cells has been functionalized *in vitro* and *in vivo* through copper-free strain-promoted alkyne-azide cycloaddition (SPAAC) to introduce a wide range of functionalities, from fluorescent dyes for tumor targeting,^33^ peptides for immunomodulation,^34^ and synthetic polymers to provide long-lasting scaffold cellularization.^35^ More recently, this chemistry has been employed for the formation of a protective layer that was found to suppress tumor growth *in vivo*,^36^ as well as for the development of living hybrid materials.^35^ These recent studies provide a relevant proof of concept for the application of MOE techniques in addressing healthcare challenges for cell-based therapeutics.

Herein, we functionalize regenerative HPC surfaces with polysaccharides, commonly used in tissue engineering applications owing to their biocompatibility and ease of modification.^37^ In particular, we explore hyaluronic acid (HA) and alginate (Alg) as cell surface functionalities, owing to their ability to establish hydrogen bonds with ECM proteins and glycans, which has been previously exploited for mucoadhesive applications.^38,39^ By covalently conjugating polysaccharide moieties through MOE and bio-orthogonal click chemistry approaches (Fig. 1a), we introduce a homogeneous, sing-cell coating that successfully increases *in vitro* HPC adhesion to biologically relevant surfaces of extra-cellular matrix (ECM) and vascular endothelial cells. Additionally, we employ a novel microfluidic flow chip to investigate cell adhesion to hepatic microtissue models coated with our clickable biomaterials. Our findings reveal that HA-coated HPCs exhibit increased engraftment to 3D microtissues, outlining a promising method for enhancing therapeutic cell adhesion in biomedical applications.

**Fig. 1.**
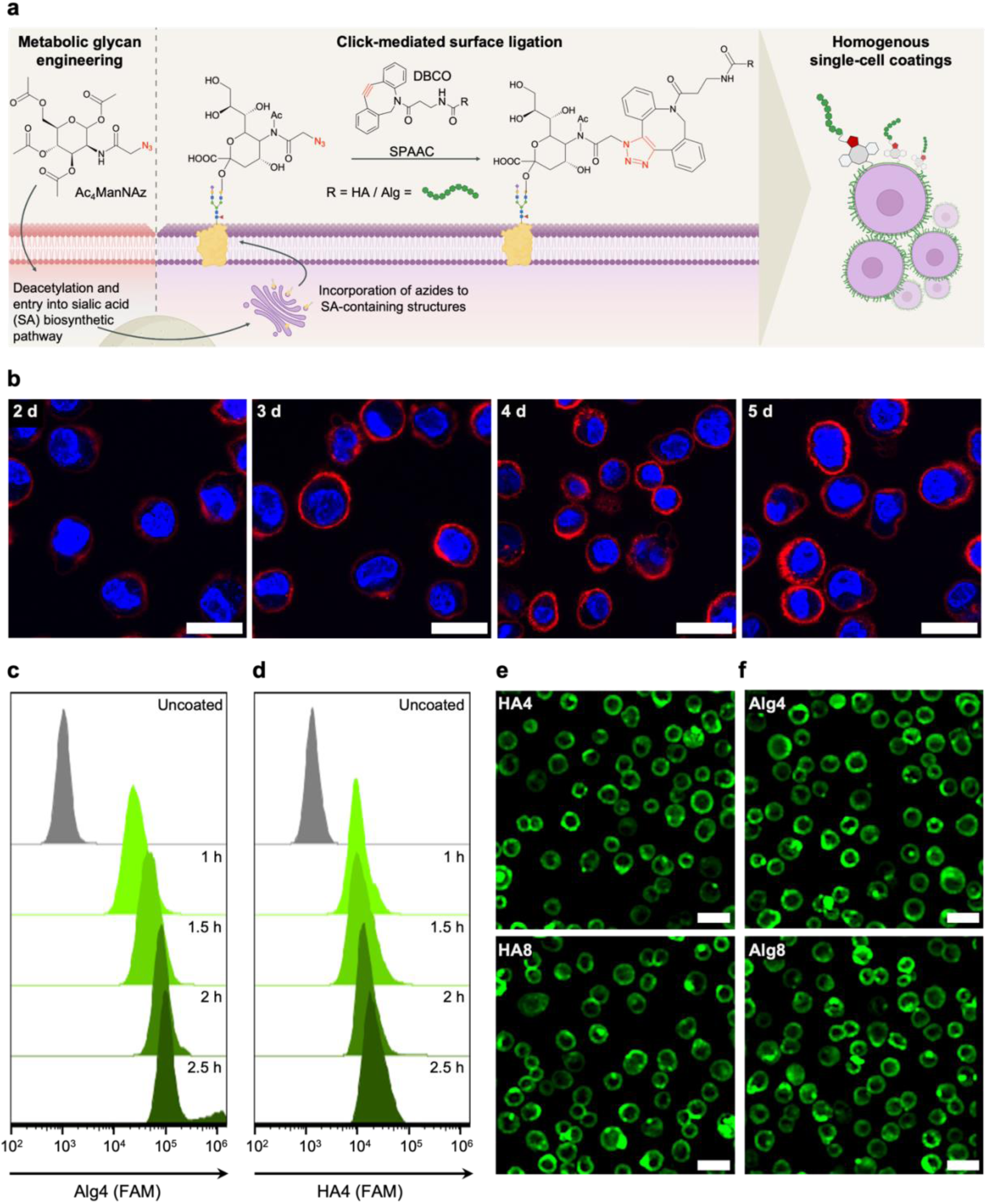
Whole-cell coating of HPCs through MOE. **a.** Schematic of metabolic glycan labelling of mammalian cells resulting in the incorporation of *N*-azidoacetyl-D-neuraminic acid (NeuAz) on sialic acid (SA)-containing residues such as *N*-linked surface glycans. Supplementation of dibenzocyclooctyne (DBCO)-modified polysaccharides results in a spontaneous strain-promoted alkyne-azide cycloaddition (SPAAC) click reaction, achieving a homogenous coating at the cell surface. **b.** Confocal fluorescence images visualizing cell surface azides *via* conjugation with Cy5-DBCO, after varying the incubation length of HPCs with Ac_4_ManNAz from 2-5 days. Scale bar = 20 μm. **c, d.** Flow cytometry graphs of Ac_4_ManNAz-treated HPCs incubated with Alg4 and HA4 for the indicated time periods (1 - 2.5 h). **e, f.** Confocal fluorescence images of HPCs treated with Ac_4_MAnNAz for 4 days and subsequently incubated with HA **(e)** and Alg **(f)** with different degrees of DBCO functionalization (4% and 8%). Scale bar = 50 μm.

## RESULTS

HA and Alg were functionalized with dibenzocyclooctyne-amine (DBCO-amine) through 4-(4,6-dimethoxy-1,3,5-triazin-2-yl)-4-methylmorpholinium (DMTMM)-mediated amidation.^40,41^ Using this method, five and ten units of DBCO were conjugated to HA and Alg to obtain HA4, HA8 and Alg4, Alg8, with 4% and 8% degrees of functionalization as confirmed by ^1^H NMR spectroscopy (Fig. S1-4). Subsequently, fluorescein-amine (FAM) was conjugated to all polysaccharide-DBCO derivatives to yield 5% functionalization, as quantified by UV-vis spectroscopy. After removal of unreacted reagents through dialysis, the final product was characterized by ^1^H NMR spectroscopy and SEC analysis, which showed the expected increase in molecular weights of HA and Alg derivatives after the cumulative conjugation of DBCO and FAM (Fig. S5-10).

HPCs were selected for this study as they possess the advantage of *ex vivo* expansion and culture in comparison to primary hepatocytes, which require cryopreservation or immediate transplantation following harvesting.^10,42^ After expansion, HPCs were incubated with 40 µM of azide-modified mannosamine (Ac_4_ManNAz) to introduce an azide functionality at the cell surface. To optimize the level of azide groups on HPC surfaces, cells were incubated with Ac_4_ManNAz for 2 to 5 days and subsequently treated with Cy5-DBCO to allow visualization and quantification of the resultant fluorescence every 24 h through confocal fluorescence microscopy and flow cytometry. These data indicated that a maximal threshold of azide display was reached after four days of HPCs incubation with Ac_4_ManNAz (Fig. 1b and Fig. S11), with fluorescence intensity not increasing further beyond this time point; hence the four days incubation time was selected for this study.

According to previous literature, *in vitro* click-mediated cell labelling has been predominantly carried out on monolayer cell cultures, under standard cell culture conditions of 37 °C, 5% CO_2_.^30,43,44^ However, to increase translatability to a cell transplantation workflow and to achieve a more homogenous (whole-cell) coating of single cells, we sought to carry out click-mediated coating on suspended HPCs. Our preliminary attempts at surface functionalization of suspended cells at 37 °C showed significant non-specific binding of Cy5-DBCO and DBCO-FAM-functionalized HA and Alg, likely caused by extensive internalization of the biopolymers and non-specific membrane interactions even in in non-azide treated cells (Fig. S12). These observations are in alignment with findings reported by Gibson and co-workers, where Cy3-DBCO and DBCO-functionalized poly(hydroxyethyl acrylamide)(pHEA)-FAM were also internalized by azide-treated human lung cancer fibroblasts.^45^ For this reason, we explored incubation at 4 °C, which is known to reduce cellular metabolic activity. Indeed, this resulted in a significant decrease in non-specific binding and internalization, for both the small molecule Cy5-DBCO and the DBCO-FAM-functionalized biopolymers (Fig. S12).

To determine the maximum viable concentration of DBCO polysaccharides to be used for coating, azide-treated HPCs were incubated with varying concentrations of HA-DBCO and Alg-DBCO (0-3 mg mL^-1^) for 2.5 h and viability monitored over 72 h using a PrestoBlue viability assay (Fig. S13). Both polysaccharides showed high cytocompatibility up to a concentration of 2 mg mL^-1^. Interestingly, HA-DBCO coated cells exhibited over 100% viability, likely as a consequence of the proliferative effect of HA, previously reported in the literature.^46,47^

To determine the optimal incubation time, HPCs were incubated at 4 °C under shaking with 1.5 mg mL^-1^ of either functionalized HA or Alg, followed by flow cytometry analysis to quantify polymer conjugation (Fig. 1c, d). As expected, a slight increase in fluorescence was observed for cells coated with HA8 and Alg8 compared to HA4 and Alg4, respectively accounting for more polymer conjugation at the cell surface with increasing DBCO functionalization (Fig. S14). Furthermore, greater fluorescence intensity was observed for both polymers when incubation time was increased from 1 h to 2.5 h (Fig. 1c, d). However, confocal fluorescence microscopy images revealed increased intracellular fluorescence after 2.5 h of incubation, likely as a result of polymer internalization over time (Fig. S15). As such, an incubation period of 2 h was preferred and adopted in all our experiments. Confocal fluorescence microscopy images of Ac_4_ManNAz-treated HPCs incubated for 2 h with HA (Fig. 1e) and Alg (Fig. 1f) derivatives demonstrate that homogeneous, whole-cell encapsulation could be achieved within this timeframe.

One critical advantage of MOE in the context of cell-based therapies is the covalent linkage of payloads throughout the cell surface. This offers the possibility to shed the polymer coating from the cell membrane in a physiological process, allowing uncoated cells to establish new interactions with the ECM and surrounding tissue *in situ*.^48^ For our HPCs, the coating started to visibly detach from the cell surface after 24 h of incubation at 37 °C, 5% CO_2_, when cells were left to adhere on a collagen coated dish. Complete disappearance of the polymer coating could be observed after 96 h, as evidenced by both confocal fluorescence microscopy (Fig. 2a, b, c, and Fig. S16) and flow cytometry (Fig. 2d), with a level of quantified fluorescence comparable to uncoated cells reached at the end of this time window. As expected, no difference was observed between the two polysaccharide coatings or among the different degrees of DBCO functionalization, considering the polymer coating shedding is solely dependent on the rate of membrane turnover specific to each cell population.

**Fig. 2.**
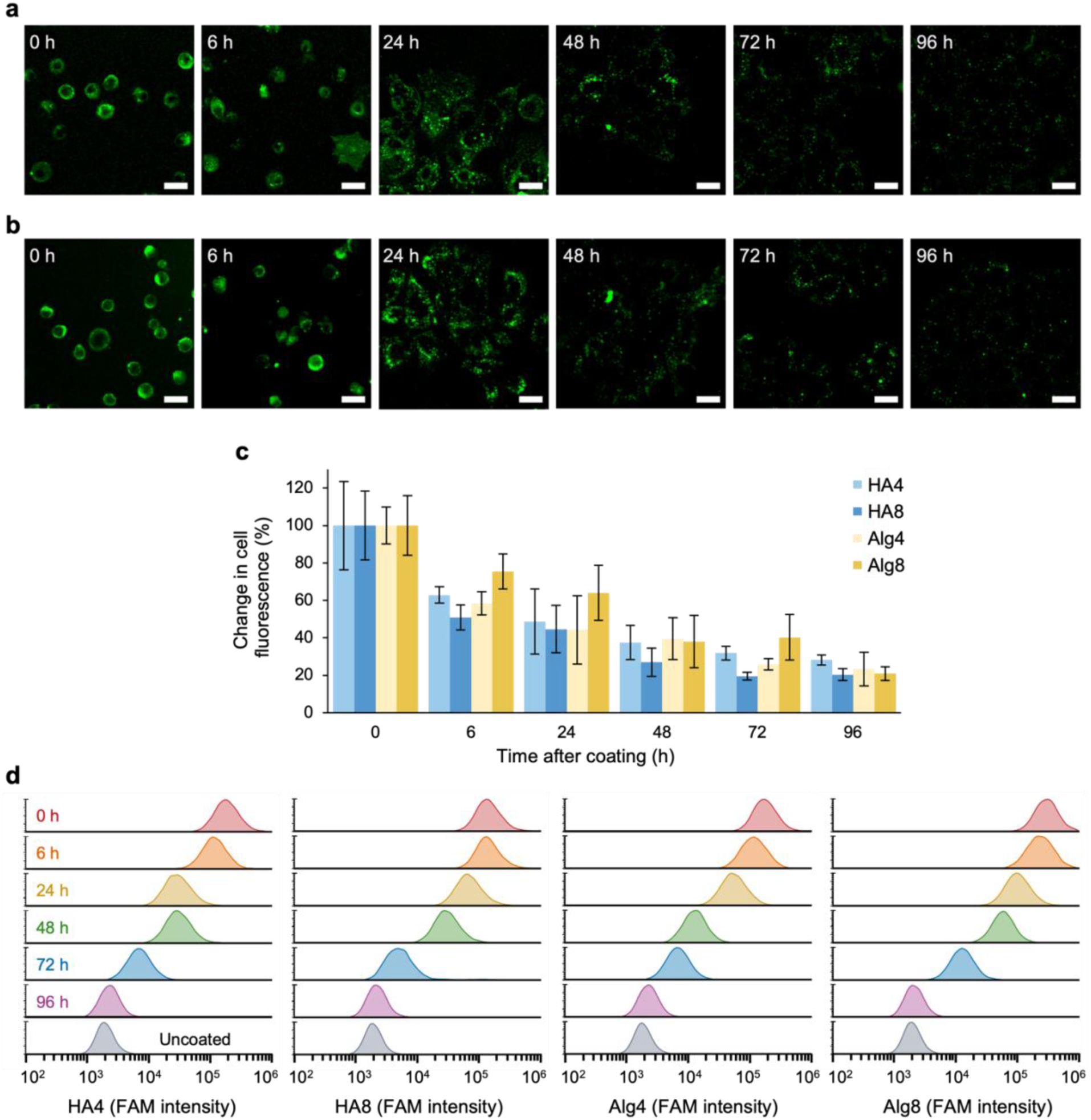
Transient surface lifetimes of conjugated polysaccharides. Confocal fluorescence microscopy images of HPCs in standard cell culture media after coating with HA4 **(a)** and Alg4 **(b)** captured over a period of 96 h. Scale bar = 20 µm. **c.** The mean relative FAM fluorescence of coated HPCs was quantified as a function of cell number per field area by analyzing captured confocal fluorescence microscopy images using FIJI (N = 3 ± SD). **d.** Flow cytometry analysis of cell surface fluorescence of HPCs coated with HA and Alg at different degrees of DBCO functionalization, showing a decrease in fluorescence over time.

The integrin family are key factors in cell binding to tissues and play a role in engraftment of regenerative cells to the liver.^49^ To characterise the potential effect of coatings on integrin expression, we assessed mRNA expression for an array of integrins (Fig. S17). Integrin expression did not decrease 20 h after coating and in the case of integrins α5, α6, α7, and β3, β5, β8 an increase in expression was observed. This could potentially play a role in facilitating and maintaining enhanced engraftment as the polysaccharides disappear from the cell surface.

In order to test our hypothesis and assess whether the polysaccharide coating was indeed able to increase cell adhesion, a colorimetric ECM adhesion assay was performed as an indirect measure of cell adhesion to an array of ECM proteins.^50^ As expected, a significant increase in adhesion to multiple ECM proteins and glycoproteins was observed for HA-DBCO derivatives, up to 92% higher compared to uncoated cells (Fig. 3a). The difference in adhesion observed between the different collagen (Col) subtypes is likely a consequence of the intrinsic differences in isoform structure which translate in different HA interactions.^51^ For example, Col-IV is commonly found in basement membranes and naturally forms sheet-like structures rather than fibrils, thus not natively binding or interacting with HA. In contrast, Col-II forms a looser, mesh-like structure which may account for increased interaction, and thus increased adhesion, in comparison to Col-I which instead forms more densely packed fibril bundles. Surprisingly, no significant increase in HPC adhesion was found for fibronectin coated surfaces, which contain specific hyaluronic acid binding motifs, suggesting that HPCs already natively bind at maximal levels to fibronectin. On the other hand, the significant increase in adhesion observed for laminin, which is found in basal lamina, is likely a consequence of the extensive post-translational glycosylation, which confers interactive hydrogen binding sites for HA compared to collagen.^52^ Similar type of secondary interactions may be attributed to the increased adhesion levels of HA-coated cells for tenascin and vitronectin.^53,54^ In contrast, HPCs coated with Alg4 and Alg8 showed significantly reduced binding to all ECM surfaces (Fig. 3a). This may be explained by the intrinsic capability of alginate to form ionic crosslinks with divalent cations such as Ca^2+^, present in the cell culture medium. This crosslinking likely results in the formation of a relatively stiffer layer at the cell surface, effectively immobilizing the cell in an encapsulating hydrogel. While hyaluronic acid is also able to crosslink with divalent cations, the presence of one carboxylic acid group per repeating unit compared to two carboxylic acid groups in the case of alginate, accounts for the lower degree of crosslinking, hence leading to weaker hydrogel coatings. To explore this, we analyzed the morphologies of HA and Alg coated cells after 2 h of seeding onto Col-I tissue culture dishes (Fig. 3c, d, and Fig. S18). FIJI image analysis of segmented cells allowed for the quantification of spreading area and cell circularity. We found HA-coated cells showed a significant increase in spreading area, with increased formation of membrane extensions, as indicated by lower values of circularity. On the other hand, alginate exhibited significantly lower cell spreading and higher values for circularity, hence supporting the hypothesis for stiff surface hydrogel formation. This is also in agreement with the increased integrin expression seen for Alg-coated cells, where a higher surface stiffness is linked with a higher level of integrin expression and cell-cell interactions (Fig. S17).^55–57^

**Fig. 3.**
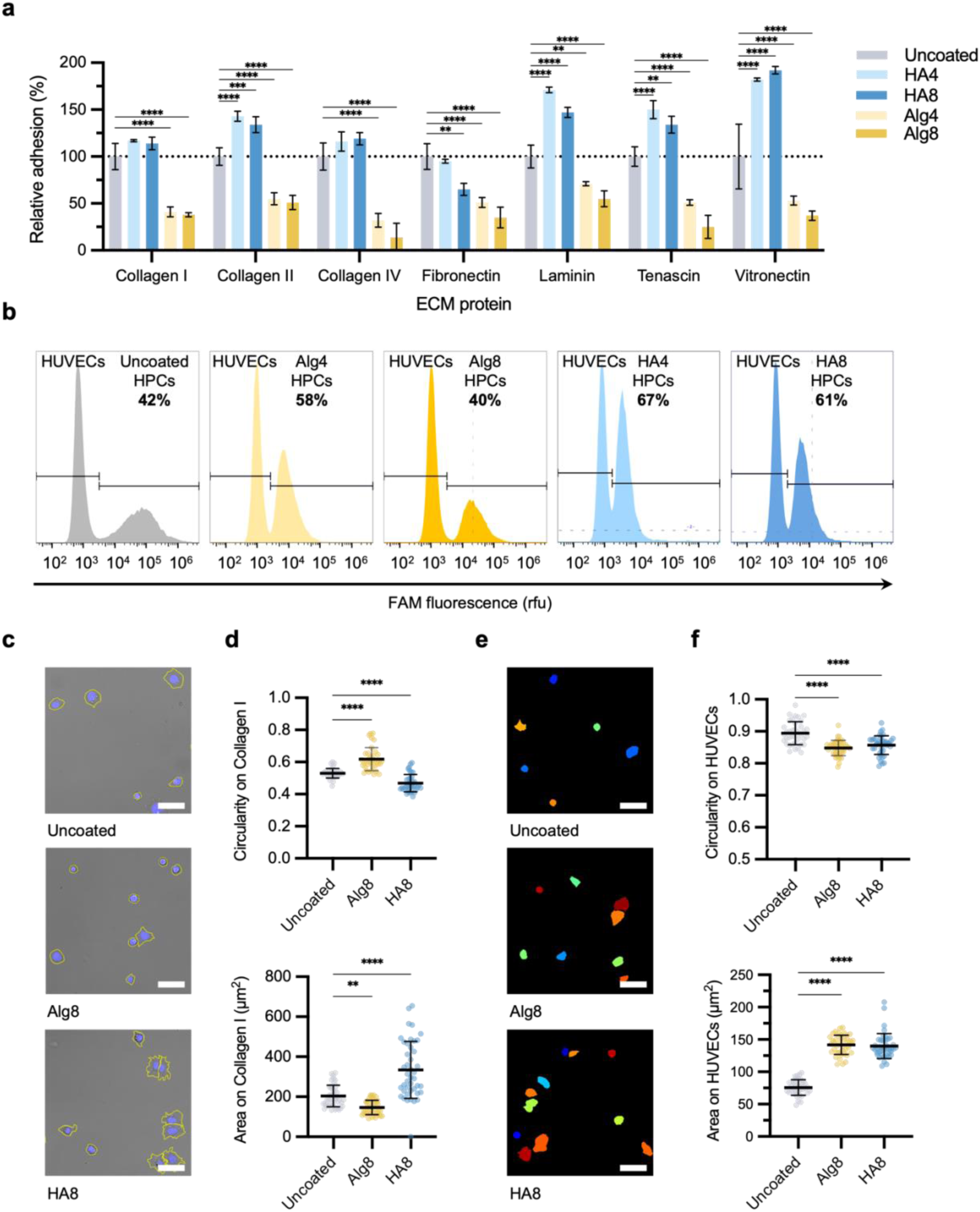
Polysaccharide coatings modulate cell adhesion to ECM proteins and human umbilical vein endothelial cell (HUVEC) monolayers. **a.** Quantification of HPCs coated with Alg and HA after adhesion onto ECM-coated 96-well surfaces, relative to uncoated HPCs. Data are shown as means of N = 3 ± SD (**** P = <0.0001). **b.** Flow cytometry graphs of uncoated HPCs labelled with green fluorescent protein or coated HPCs after adhesion onto HUVEC monolayers for 2 h. Unbound cells were removed by PBS washings and the relative proportions of HPCs adhered to HUVECs were quantified by flow cytometry after subsequent dissociation with accutase. **c.** Confocal brightfield images of uncoated, Alg8, and HA8 coated HPCs left to adhere for 2 h on Col-I coated dishes. Yellow outlines indicate cell perimeters. Scale bar = 50 μm. **d.** Morphological parameters for area and circularity analyzed from confocal brightfield images for uncoated, Alg8, and HA8 coated HPCs after adhesion onto collagen-coated plates (N = 3 ± SD, **** P = <0.0001). **e.** Confocal brightfield image masks of uncoated, Alg8, and HA8 coated HPCs left to adhere for 2 h on HUVEC monolayers. Scale bar = 50 μm. **f.** Morphological parameters for area and circularity analyzed from confocal brightfield image masks for uncoated, Alg8, and HA8 coated HPCs after adhesion onto HUVEC monolayers (N = 3 ± SD, **** P = <0.0001).

In order to determine the ability of coated HPCs to still interact and adhere to other cells, we explored their behavior on confluent monolayers of human umbilical vein endothelial cells (HUVECs) as a model for vasculature endothelium interactions. HUVECs were seeded onto collagen-coated plates to model cell-cell interactions of HPCs to the sinusoidal microenvironment of the liver. Uncoated HPCs, fluorescently labelled with green fluorescence protein (GFP), as well as HPCs coated with Alg or HA were seeded on top of HUVEC monolayers and left to adhere for 2 h. Following extensive PBS washing and dissociation, flow cytometry analysis was used to quantify the relative number of bound cells which formed tight cell-cell focal adhesions (Fig. 3b). HPCs coated with HA4 exhibited the highest level of adhesion to the HUVEC monolayer, with a measured ratio of HPCs to HUVECs of 67%. Similarly, HPCs coated with HA8 also showed strong adhesion, with a measured ratio between the two cell types of 61%. Interestingly, HPCs functionalized with Alg8 also showed increased adhesion compared to uncoated control, with a population value of 58%. This was surprising, considering the previous results obtained with our ECM adhesion model, however it could be explained with the oversimplified nature of ECM adhesion assays, which do not take into account the active participation of partner cells in cell-cell interactions. Consistently with these results, quantification of morphological parameters, including circularity and cell area suggest that HPCs coated with both HA8 and Alg8 are able to spread better on HUVEC monolayers compared to uncoated HPCs, indicating a higher level of cell-cell interactions and adhesion (Fig. 3e, f, and Fig. S19).

To further expand the complexity of our adhesion models, we sought to introduce flow and shear as a physiologically relevant parameter to our adhesion assays. Akura^TM^ ImmuneFlow chips were used to study adhesion to 3D InSight™ human liver microtissues (hLMTs), mimicking the *in vivo* environment where cells travel through blood vessels before reaching the site of engraftment. Moreover, the hLMTs are constructed with a range of different cells, including human hepatocytes, Kupfer cells, and lymphatic endothelial cells, mimicking the hepatic microenvironment found *in vivo*. In this microfluidic platform HPCs remain in suspension and continuous flow, using gravity and repeated back and forth tilting of the chip (Fig. 4a). Adhesion of HPCs is tested while they pass the 3D microtissues loaded in special culturing compartments, rather than static 2D cell monolayers. Considering HA coating has shown the most promising adhesion to both ECM components and HUVEC monolayers, we sought to explore both HA4 and HA8 coated HPCs for their adhesion to hLMTs. Confocal fluorescence microscopy analysis of HPCs coated with HA4 and HA8 showed increased adhesion to the liver microtissues compared to uncoated HPCs (Fig. 4b). Moreover, quantification by flow cytometry showed a substantial increase in adhesion for HA4 and HA8 coated HPCs when compared to uncoated cells (3.2 and 3.4 fold, respectively) after 9 h of incubation under flow conditions (Fig. 4c). This demonstrates that our technology platform can be successfully translated in a setting that mimics an *in vivo* environment.

**Fig. 4.**
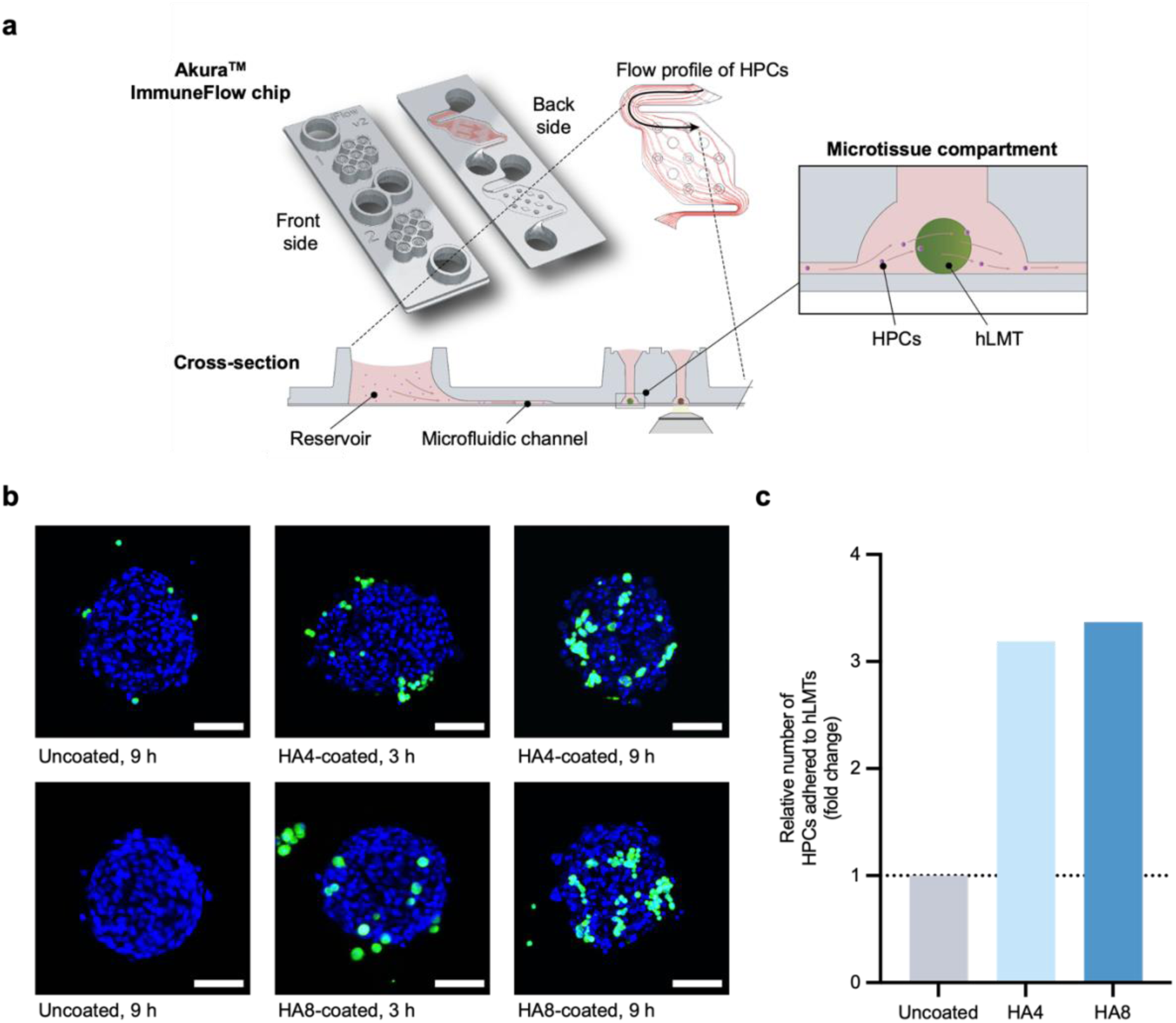
Engraftment of coated HPCs to liver microtissues under microfluidic flow. **a.** Schematic representation of InSphero Akura^TM^ ImmuneFlow chip used for the study of coated HPCs engraftment to hLMTs under gravity-induced flow. **b.** Representative confocal fluorescence images of uncoated and HA-functionalized HPCs engrafted to hLMTs after 3 h and 9 h of incubation under flow (N = 7). Uncoated and coated cells were stained with CellTracker deep-red before injection in the microfluidic channels. Scale bar = 100 μm. **c.** Cumulative flow cytometry quantification of uncoated, HA4, and HA8 coated HPCs engrafted to hLMTs after 9 h of incubation in flow. Data are presented relative to uncoated HPCs.

## DISCUSSION

Cell-based regenerative therapies for liver disease are currently hindered by suboptimal efficacies as a result of low engraftment post-transplantation. The heterogenous surface of the cell membrane, controlling major cellular functions of cell-cell communication and environmental sensing, represents a rational target for modification to enable manipulation of cell adhesion mechanisms. Traditional genetic modification approaches of cell surface proteins and associated enzymes are widely used in cell engineering. Indeed, it has been shown that the genetic overexpression of HA synthases significantly increases the production of HA in cellular models, directly resulting in increased adhesion to diverse surfaces and driving the formation of membrane extensions.^58–60^ In this work, we propose the use of MOE for the click-enabled attachment of adhesive polysaccharides to the cell surface, with the aim to increase regenerative efficacies of HPCs by increasing their adhesive capacities to ECM and surrounding tissue, circumventing the associated drawbacks of genetic modification methodologies and allowing to reduce preparation time. Moreover, the transient nature of the conjugated polysaccharide functionalities aligns well with the regenerative timescale, as sustained motifs enhancing cell adhesion become redundant post-engraftment at targeted sites. Upon transplantation into the diseased liver microenvironment, the enhanced adhesion capacity of HA-coated HPCs to ECM proteins of sinusoidal endothelial cells serves as a pivotal initial interaction, facilitating subsequent integrin-mediated adhesion and engraftment. Notably, our investigations extend to showcasing enhanced adhesion to 3D microtissue models of liver parenchyma, mimicking the situation coated HPCs encounter *in vivo*.

The translatability of this strategy opens the door for the improvement of cell-based therapies and further explorations of adhesive macromolecules for the enhanced engraftment of regenerative cells.

## METHODS

### HA and Alg functionalization with DBCO and FAM

500 mg (11.1 µmol) of low-MW HA (Biosynth, FH01773) was dissolved in 10 mL of water with 8 eq. (24.6 mg, 88.8 µmol) of DMTMM (4-(4,6-Dimethoxy-1,3,5-triazin-2-yl)-4-methylmorpholinium chloride). 6 eq. (18.4 mg, 66.6 µmol) of dibenzocyclooctyne-amine (Sigma-Aldrich) dissolved in a 5 mL volume of tetrahydrofuran was slowly added to the solution and the reaction mixture was left to stir for 5 days to yield HA4. Molar equivalents were doubled to yield HA8. The product was then purified through dialysis for 3 days (MWCO 3.5 kDa) and lyophilized to yield a white product (87% and 93% yields for HA4 and H8, respectively). Alg4 and Alg8 were functionalized in a similar fashion, from a starting quantity of 500 mg (9.09 µmol) of low-MW Alginate (FMC biopolymers, Protanal LFR 5/60), yielding 85% and 88%, respectively. A 10 mg sample of DBCO-functionalized HA or Alg was subjected to a solubilization/lyophilization cycle in deuterium oxide (D_2_O), three times. It was then redissolved in D_2_O for ^1^H NMR spectroscopy analysis. Spectra were recorded on a Bruker 400 MHz spectrometer at 298 K and analyzed using MestReNova software. Each polysaccharide-DBCO variant was functionalized with FAM by dissolving 200 mg of each polysaccharide variant (4.3µmol, 4.2µmol, 3.6µmol, and 3.5µmol for HA4, HA8, Alg4, Alg8, respectively) in 15 mL of 50:50 H_2_O:EtOH, followed by the addition of 6 eq. of FAM (9 mg, 25.8 µmol for HA4, 8.8 mg, 25.2 µmol for HA8, 7.5 mg, 21.6 µmol for Alg4, and 7.3 mg, 21 µmol for Alg8). FAM quantification was performed by obtaining a UV-vis standard calibration curve of FAM at 488nm in a 50:50 mixture of water and ethanol. The UV-vis absorption of 0.05% w/v of FAM-functionalized polysaccharides was measured, and FAM content of the solution was subsequently extrapolated using Beer’s law.

### Cell surface functionalization and analysis

Prior to incubation with the polysaccharide solution, fresh HPCs were seeded into fresh Col-1 treated T75 flasks at a cell density of 3 × 10^5^ cells per T75 cell culture flask. Standard DMEM solution supplemented with 40 µM Ac_4_ManNAz was added to cells, which were then left to proliferate for four days prior to coating at 37 °C in a humidified atmosphere containing 5% CO_2_. DBCO-FAM functionalized HA or Alg was then dissolved in PBS, with the aid of gentle heating and a vortex mixer to obtain a 0.75% w/v stock solution. HPCs were detached using accutase cell dissociation solution to yield a cell suspension in DMEM. 1.5 × 10^6^ cells were resuspended 1.8 mL DMEM supplemented with a final concentration of 0.15% w/v from the polysaccharide stock solution. Cells were left to incubate in a cooled shaking incubator for 2 h (4 °C, 80 rpm), followed by washing with PBS (3 × 1 mL) and filtration using 40 µm cell strainers (3×). Cell suspensions were then either placed in 35 mm confocal dishes for confocal fluorescence microscopy analysis, adhesion analysis or flow cytometry characterization.

### Viability assays and quantification of HA and Alg lifetimes

Viability of cells incubated with non-fluorescent DBCO-functionalized polysaccharides was quantified by seeding cells onto Col-I coated 24-well plates seeded with 3.8 × 10^4^ cells per well, followed by the addition of PrestoBlue cell viability reagent (Invitrogen, A13261) at each time point (24 - 72 h), following manufacturer’s protocol. Endpoint fluorescence measurements at 590 nm were taken with a FLUOstar Omega microplate reader (BMG Labtech) using Omega MARS software (BMG Labtech).

The lifetime of DBCO-FAM polysaccharides coating on HPCs was quantified by seeding the cells on Col-I treated 35 mm confocal dishes with 1 × 10^5^ coated or uncoated cells. At each time point considered (6 - 96 h), HPCs were washed with PBS (3 × 1 mL) followed by live cell confocal fluorescence microscopy imaging. At least five Z-stack images (imaging depth of 20 µm) from each dish (N = 3) were captured at each time point to monitor the presence of HA or Alg on the cell surface. Average cell fluorescence intensities were quantified using Olympus CellSens software in which a set value for image thresholds were applied to all images in the FAM channel followed by ROI selection of cells and fluorescence intensity quantification. Fluorescence intensities of coated cells were also quantified by flow cytometry at each time point, following treatment with accutase cell dissociation reagent.

### Quantification of HPC adhesion to ECM and HUVECs

HPCs were coated with non-fluorescent DBCO-functionalized HA or Alg as outlined above and seeded on ECM540 adhesion array 96-well plate kit (1 × 10^5^ cells), following manufacturer’s protocol (Merck). Cells were incubated for 1 h at 37 °C, 5% CO_2_. After incubation, media was discarded and cells were washed with assay buffer (3 × 100 µL), followed by 5 min incubation at RT with cell stain solution and washed with PBS (3 × 100 µL). Stained cells were left to air dry for 5 min, followed by treatment with extraction buffer and gentle shaking on an orbital shaker for 10 min. Endpoint absorbance at 544 nm was taken with a FLUOstar Omega microplate reader (BMG Labtech) using Omega MARS software (BMG Labtech).

HUVECs (Gibco, C01510C) were cultured as standard in Human Large Vessel Endothelial Cell Basal Medium (Gibco, M200500) supplemented with low serum growth supplement (Gibco, S00310). Cell maintenance was carried out by replacement of medium every two days and passaging upon reaching around 60% confluency by dissociation with trypsin-EDTA solution. To obtain HUVEC monolayers, cells were seeded on either 6-well plates or 35 mm confocal microscope dishes pre-coated with Col-I at a cell density of 5 × 10^4^ cells cm^-2^. At 100% HUVEC confluency, 1 × 10^6^ coated or uncoated GFP^+^ HPCs were seeded onto the HUVEC monolayers and left to incubate for 2 h, followed by washing with PBS (3 × 1 mL). The resulting co-culture of cells was then analyzed by flow cytometry. Flow cytometry samples required prior dissociation into a single cell suspension by treatment with accutase cell dissociation reagent and resuspension in PBS flow buffer supplemented with 3% v/v FBS and 3% v/v EDTA. Cells were then filtered using a 40 µm cell strainer.

### Cell morphology characterization

HPCs were coated with HA or Alg as described above and seeded onto collagen-coated 35 mm confocal dishes at a seeding density of 5 × 10^5^ cells per dish. Cells were left to adhere for 2 h and then washed with PBS (3 × 1 mL) to remove non-adherent cells. Nuclei were then stained with 1 µM Hoechst 33342 for 15 min. Confocal laser scanning microscopy was used to capture a minimum of 12 images from each dish. Morphological parameters were quantified by outlining cell perimeters as regions of interest and using the measure feature in the brightfield. At least 700 cells per experiment were measured.

For morphological analysis of cells adhered to HUVECs, 35 mm confocal microscope dishes containing 100% confluent monolayers of HUVECs were prepared as described before. Prior to seeding, HPCs (coated or uncoated) were stained with Celltracker DeepRed dye, following manufacturer protocol (1 µM, 10 min). 5 × 10^5^ cells were seeded per dish and left to adhere for 2 h, after which, unbound cells were washed with PBS (3 × 1 mL). Nuclei were stained with 1 µM Hoechst 33342 for 15 min. A minimum of 12 confocal laser scanning microscope images were then captured from each dish followed by CellProfiler analysis of the Cy5 fluorescence channel to quantify HPC morphology.

### Adhesion assays using Akura^TM^ ImmuneFlow chips

Akura^TM^ ImmuneFlow chips (https://insphero.com/3d-cell-culture-tools/akura-immune-flow-organ-on-chip-platform/) were prepared according to manufacturer’s protocol. hLMTs (InSphero, MT-02-302-04) were loaded into the microtissue compartment, followed by the addition of uncoated and coated HPCs (labelled with CellTracker deep-red) in a 200 µL DMEM suspension at a cell density of 50 × 10^3^ cells mL^-1^. The microfluidic chips were loaded onto Akura™ All-in-one Tilting System (InSphero) following a tilting program of: ± 85° tilt (25 s movement, 1:15 min hold), horizontal pause (5 min).

hLMTs were imaged at time points of 3 h and 9 h. At the 9 h time point, PBS was flowed through the chips and hLMTs were collected and dissociated by the addition of 200 µL accutase with gentle vortexing for 30 min. The dissociation of hLMTs was routinely monitored using an inverted light microscope. Following complete dissociation, single cell suspensions were analyzed by flow cytometry quantifying for CellTracker deep-red labelled HPCs.

## Acknowledgements

A.R.A. thanks EPSRC and SFI Centre for Doctoral Training in Engineered Tissues for Discovery, Industry and Medicine (grant number EP/ SO2347X/1) for support through a Ph.D. scholarship. UK Research and Innovation (UKRI, Future Leaders Fellowship, MR/X033546/1) are thanked for supporting M.C.A. MRC UKRMP ECE Hub (# 623924) and MRC Exploiting In Silico Modelling To Address The Translational Bottleneck In Regenerative Medicine Safety (# 996932) are thanked for support to A.J.E.H. and M.R.V. ERC Advanced Award #789119 is thanked for support to A.J.E.H. HPCs were provided as a kind gift from Dr Wei-Yu Lu (University of Edinburgh). GPF-labelled HPCs were kindly donated by Dr Candice Ashmore-Harris and Prof. Stuart Forbes (University of Edinburgh). Figure 1a was created using Biorender.com.

## Author contributions

M.C.A. and A.J.E.H. designed the study. T.H., L.H., and O.F. designed the AkuraTM ImmuneFlow chip and trained A.R.A. in performing the experiments. A.R.A. performed the majority of the experiments and data analysis. M.R.V. performed the PCR experiments and conducted data analysis on those. A.R.A., M.C.A. and A.J.E.H. wrote the manuscript with the contribution of all authors.

## Supplementary information

Supplementary methods and figures are available in the supplementary information file.

