## Supplementary information for "Sweet and sticky: increased cell adhesion through click-mediated functionalization of regenerative liver progenitor cells"

**NMR spectroscopy.** $^1$H NMR spectra were recorded at 400 MHz on a Bruker DPX-400 spectrometer in $D_2O$, unless otherwise stated. Chemical shifts are reported as $\delta$ in parts per million (ppm) downfield from the internal standard trimethylsilane.
NMR spectroscopy assignments for Alg-DBCO: $^1$H NMR (400 MHz, $D_2O$) $\delta$/ppm: 7.71-7.40 (8H, m, $C_6H_6$ from DBCO), 5.06 (1H, s, C*H* from Alg), 4.51-3.48 (6H, m, C*H* from Alg), 2.68 (1H, s, N*H*); for Alg-DBCO-FAM: $^1$H NMR (400 MHz, $D_2O$) $\delta$/ppm: 8.81-6.88 ((8H, m, $C_6H_6$ from DBCO) (9H, m, $C_6H_6$ from FAM)), 5.07 (1H, s, C*H* from Alg), 4.50-3.50 (6H, m, C*H* from Alg); for HA-DBCO: $^1$H NMR (400 MHz, $D_2O$) $\delta$/ppm: 7.72-7.38 ((8H, m, $C_6H_6$ from DBCO), 4.58-3.36 (8H, m, C*H* from HA), 2.68 (1H, s, N*H*), 2.02 (3H, m, $CH_3$ from HA); for HA-DBCO-FAM: $^1$H NMR (400 MHz, $D_2O$) $\delta$/ppm: 7.72-7.38 ((8H, m, $C_6H_6$ from DBCO), (9H, m, $C_6H_6$ from FAM), 4.58-3.36 (8 H, m, C*H* from HA), 2.02 (3H, m, $CH_3$ from HA).

**Size exclusion chromatography.** Aqueous SEC measurements were performed on an Agilent 1260 Infinity II Multi-Detector GPC/SEC System fitted with RI and ultraviolet (UV, $\lambda$ = 309 nm) detectors, using a 80:20 $H_2O$:MeOH elution solvent containing 0.1 M $NaNO_3$. Polymers were eluted though an Agilent guard column (PLGel 5 μM, 50 × 7.5 mm) and two Agilent mixed-C columns (PL aquagel-OH MIXED-H 8 μM, 300 × 7.5 mm) with a flow rate of 1 mL $min^{-1}$, 40 °C. $M_n$, $M_w$ and $Đ_M$ were determined using Agilent GPC/SEC software (vA.02.01) against a 15-point calibration curve ($M_p$ = 615 - 3,187,000 g $mol^{-1}$) based on PEG standards (Easivial PEG-M/H, Agilent).

**Confocal fluorescence microscopy.** All cell images were captured by live cell confocal microscopy using Olympus FLUOVIEW Spectral FV3000 laser scanning microscope equipped with 405, 488, 561, and 640 nm lasers. For live cell experiments, cells were kept under controlled conditions of 37 °C, 5% $CO_2$.

**Flow cytometry.** To quantify click-mediated surface functionality, flow cytometry of suspended cells was performed on a Beckman Coulter CytoFLEX flow cytometer with 4-lasers capable of 15 parameter analysis including FSC and SSC. Sample analysis required the use of the 488 nm excitation laser and a 530 nm filter for fluorescein measurements and 638 nm excitation laser and 660 filter for Cy5/Celltracker deep red measurements. All sample measurements consisted of a minimum of 30,000 total recorded events. Cells were suspended in PBS solution supplemented with 3% fetal bovine serum and 3 mM EDTA and passed through a 40 μm cell strainer to ensure single cell analysis. Voltage settings applied ensured that untreated control cells appeared at low fluorescence emission intensities (FITC and Cy5 channels) and to ensure fluorescence measurements were within the detection range ($< 10^6$ A.U.). CytExpert software (Beckman Coulter) was used for data collection. FlowJo and the online tool Floreada.io were used for data presentation.

**HPCs culture and general maintenance.** HPCs were provided as a kind gift from Dr Wei-Yu Lu (University of Edinburgh). A step-by step protocol describing the method for primary cell harvest, isolation and maintenance has been previously described.[1] Briefly, T75 flasks were coated overnight with the addition of 5 mL of 200 μg $mL^{-1}$ Col-I (Sigma-Aldrich, C3867). The Col-I solution was removed and washed with fresh PBS before use. General maintenance of the cell line was completed by passaging every 5-7 days or before reaching 90% confluency. Cells were dissociated using accutase cell dissociation reagent and re-seeded at a density of $3 \times 10^5$ cells per T75 cell culture flask. HPCs were cultured in DMEM supplemented with 10% FBS, 2.0 mM L-glutamine, and antibiotic solution containing penicillin (100 units $mL^{-1}$) and streptomycin (100 μg $mL^{-1}$) at 37 °C in a humidified atmosphere containing 5% $CO_2$. Cells were counted by standard trypan blue exclusion using an automatic cell counter (Countess II, ThermoFisher).

**RNA isolation, cDNA synthesis and qPCR**

Coated cells were seeded onto collagen-coated plates with culture media and maintained in the incubator at 37 °C and 5% $CO_2$. After 20 h in culture, cells were detached using 0.5% Trypsin-EDTA (Gibco) followed by media to inhibit the trypsin and harvested by centrifugation for 5 min. at 200 g. Cells were resuspended in TRIzol Reagent (Invitrogen) immediately after collection and stored at -20 °C before total RNA extraction. Extraction of total RNA was performed according to the TRIzol manufacturer's protocol. RNA was diluted with nuclease-free water, digested with DNase I (Invitrogen) and then stored at -80 °C until further use. RNA concentration of each sample was assessed using Spark spectrometer and NanoQuant plate (TECAN). Subsequently, total RNA was reverse transcribed to synthesise complementary DNA using SuperScript IV Reverse Transcriptase (Invitrogen), according to manufacturer's protocol. Real-time quantitative PCR was performed using SYBR Green PCR Master Mix (Applied Biosystems) according to the manufacturer's instructions on a AriaMx instrument (Agilent). PCR conditions used were as follows: initial polymerase activation at 95 °C for 15 min., followed by 40 cycles of denaturation at 94 °C for 15 seconds, annealing at 55 °C for 30 seconds, and extension at 72 °C for 30 seconds. The following commercial QuantiTect primers (QIAGEN) were used: Gapdh (QT01658692), integrin alpha 1 (QT01198554), integrin alpha 2 (QT01540798), integrin alpha 3 (QT00125678), integrin alpha 5 (QT00114611), integrin alpha 6 (QT00144354), integrin alpha 7 (QT00136990), integrin alpha 9 (QT00172459), integrin alpha v (QT00095235), integrin beta 1 (QT00155855), integrin beta 3 (QT00128849), integrin beta 4 (QT00269010), integrin beta 5 (QT00108976), integrin beta 6 (QT00128233), integrin beta 8 (QT00280686). Gene expression data was based on Cq values and quantified using the relative quantification 2-ΔΔCt method. Gene expression levels of the target transcripts were normalised against housekeeping gene GAPDH. Results were reported as mean of fold change relative to uncoated HPCs.

**Statistical analysis.** Group differences were examined with GraphPad Prism8. Statistical comparisons of multiple samples were performed using a one-way analysis of variance (ANOVA), followed by Dunnett's multiple comparisons test. A P value of less than 0.05 was considered statistically significant.

### Supplementary figures

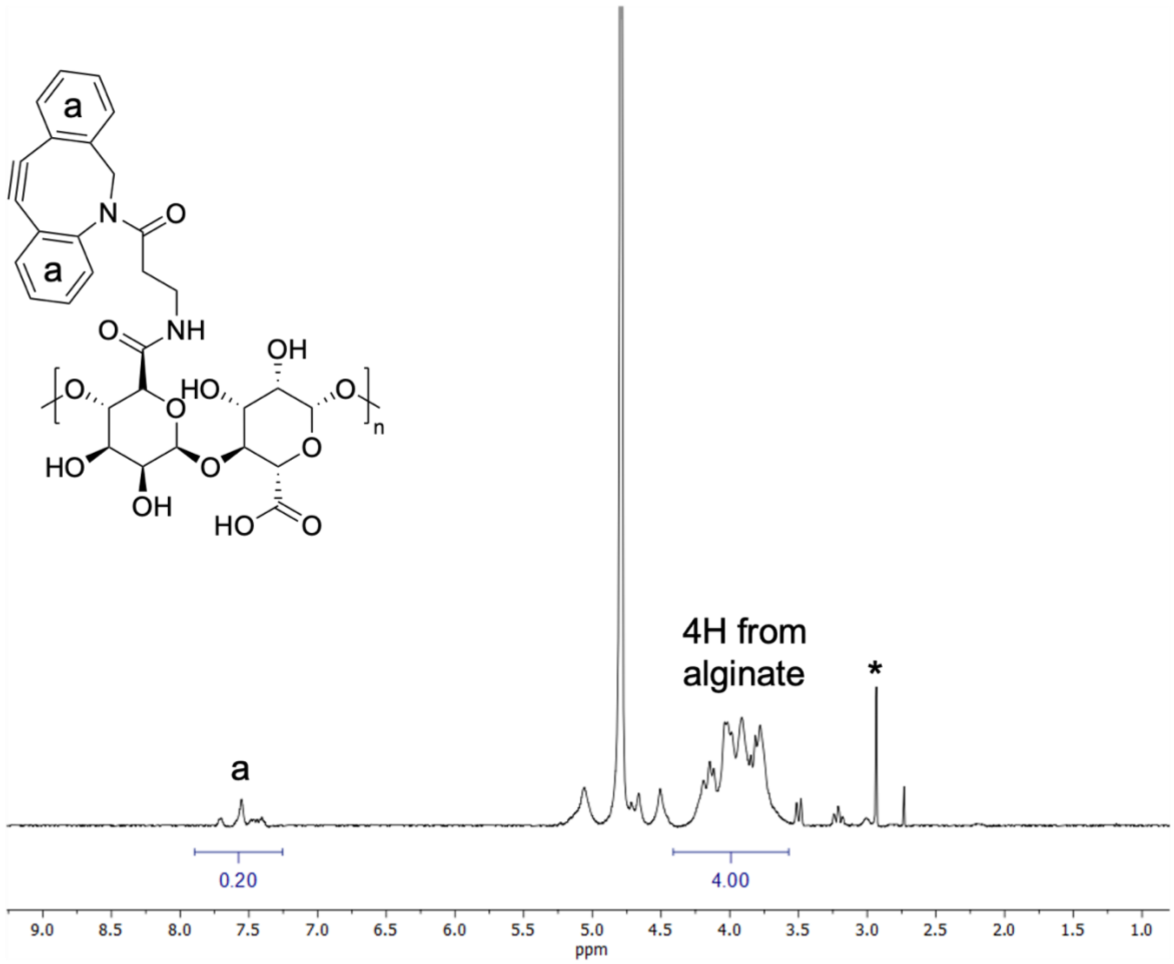


**Figure S1.** [1]H NMR spectrum (400 MHz, $D_2O$) of alginate functionalized with DBCO (Alg4). * = DMSO. Integration of the aromatic region for DBCO protons indicates a 4% functionalization (0.20 proton/8 = 0.025 units of DBCO per repeating unit, hence 5 in total).

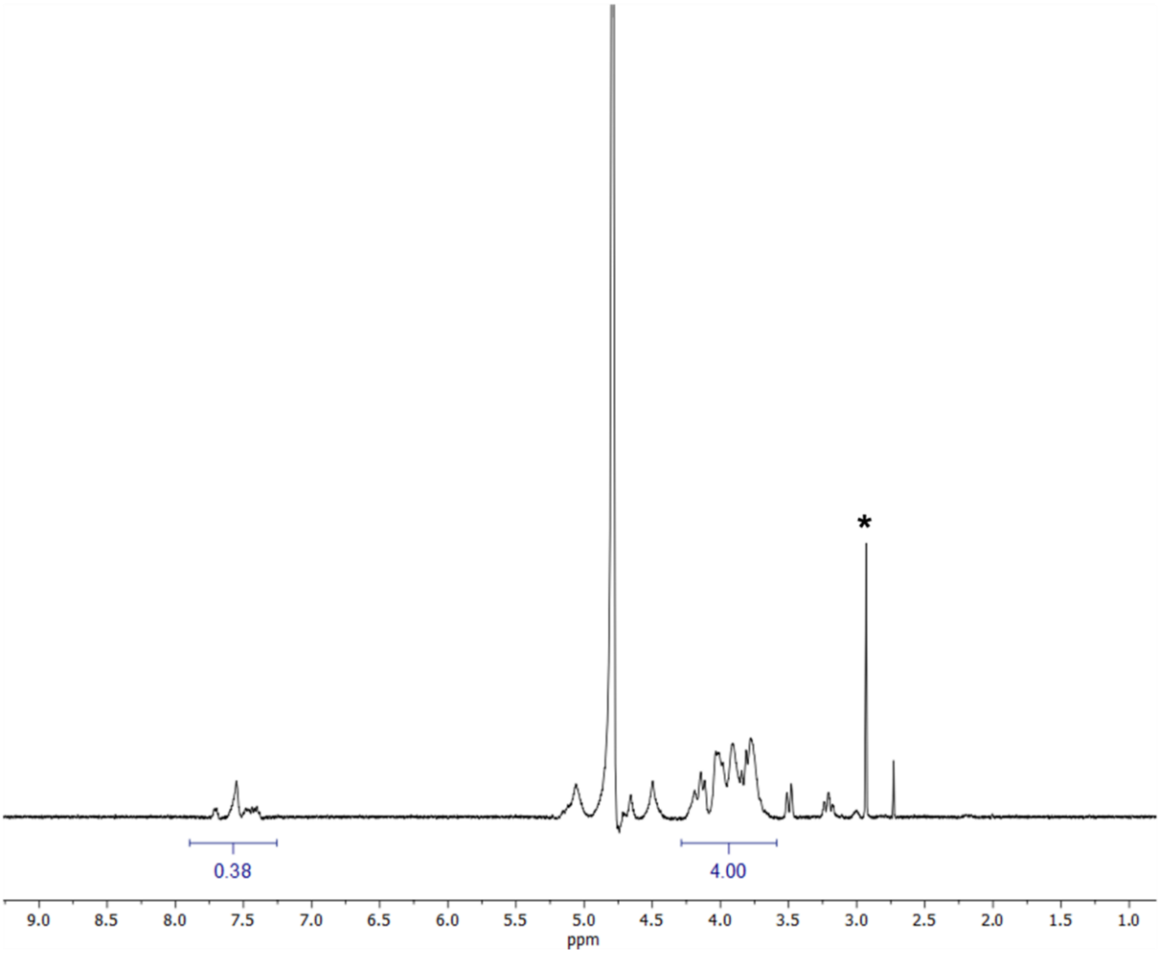


**Figure S2.** [1]H NMR spectrum (400 MHz, $D_2O$) of alginate functionalized with DBCO (Alg8). * = DMSO. Integration of the aromatic region for DBCO protons indicates an 8% functionalization (0.38 proton/8 = 0.05 units of DBCO per repeating unit, hence 10 in total).

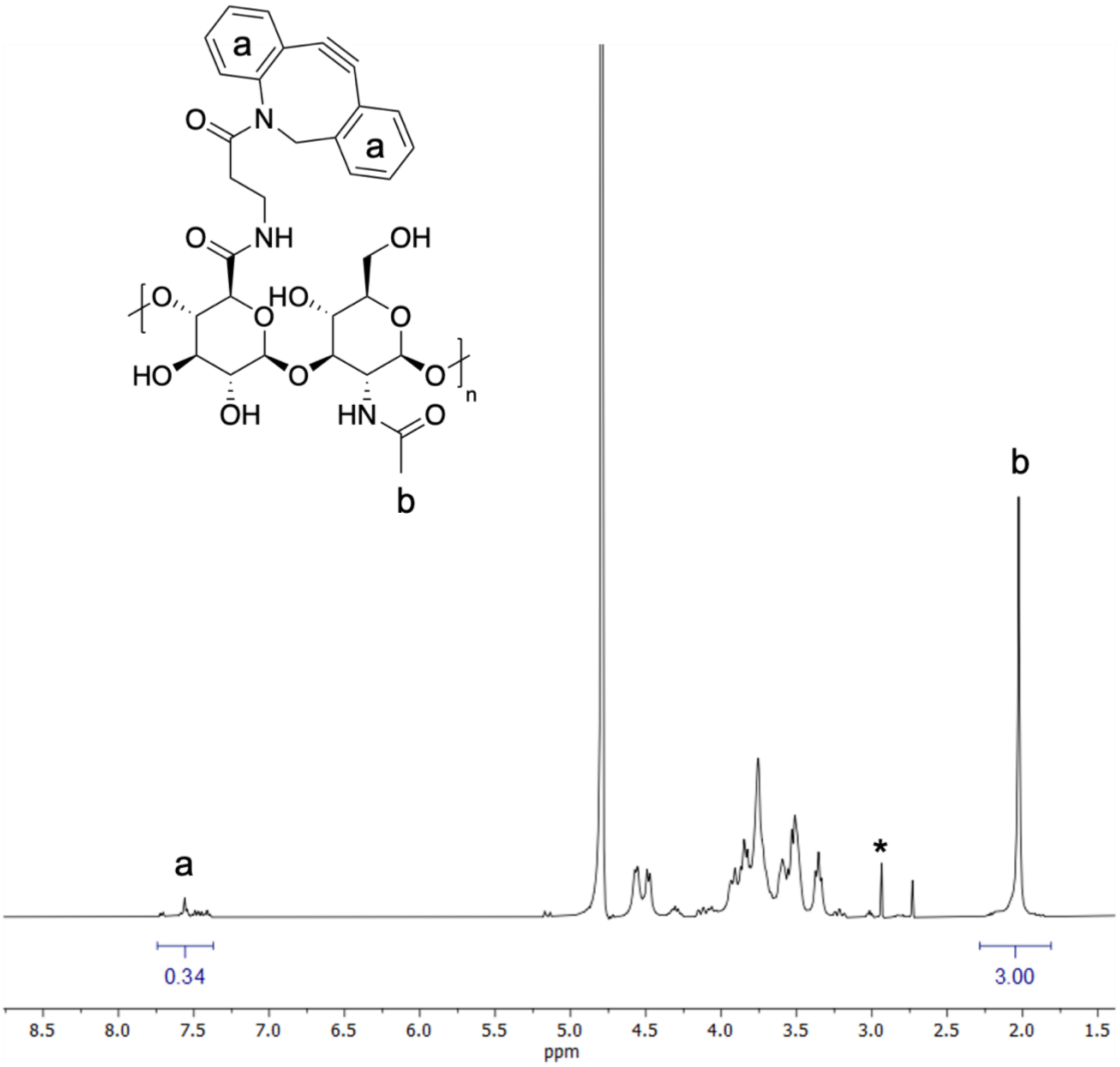


**Figure S3.** $^{1}$H NMR spectrum (400 MHz, $D_2O$) of hyaluronic acid functionalized with DBCO (HA4). * = DMSO. Integration of the aromatic region for DBCO protons indicates a 4% functionalization (0.34 proton/8 = 0.04 units of DBCO per repeating unit, hence 5 in total).

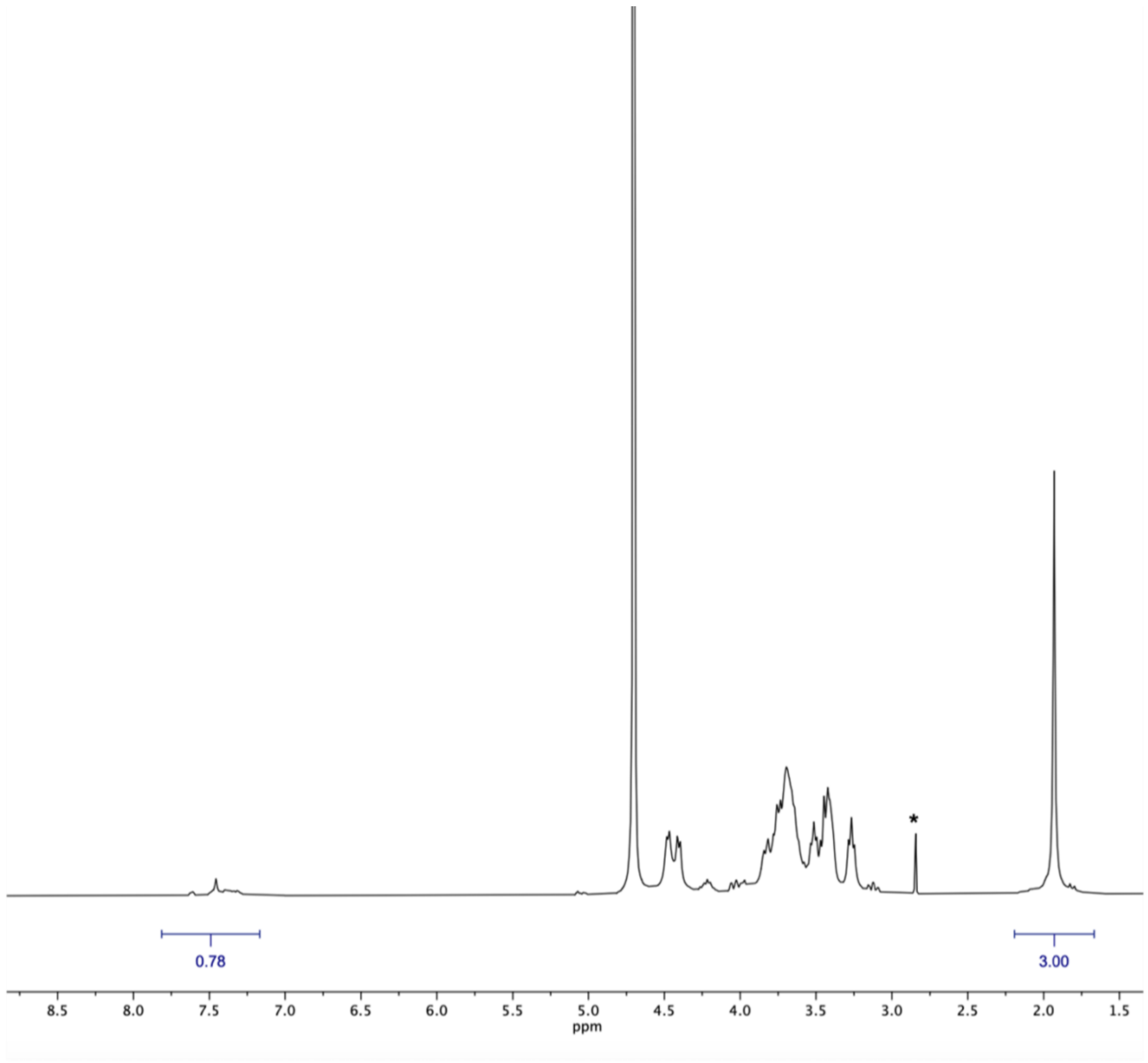


**Figure S4.** $^1$H NMR spectrum (400 MHz, $D_2O$) of hyaluronic acid functionalized with DBCO (HA8). * = DMSO. Integration of the aromatic region for DBCO protons indicates an 8% functionalization (0.78 proton/8 = 0.1 units of DBCO per repeating unit, hence 11 in total).

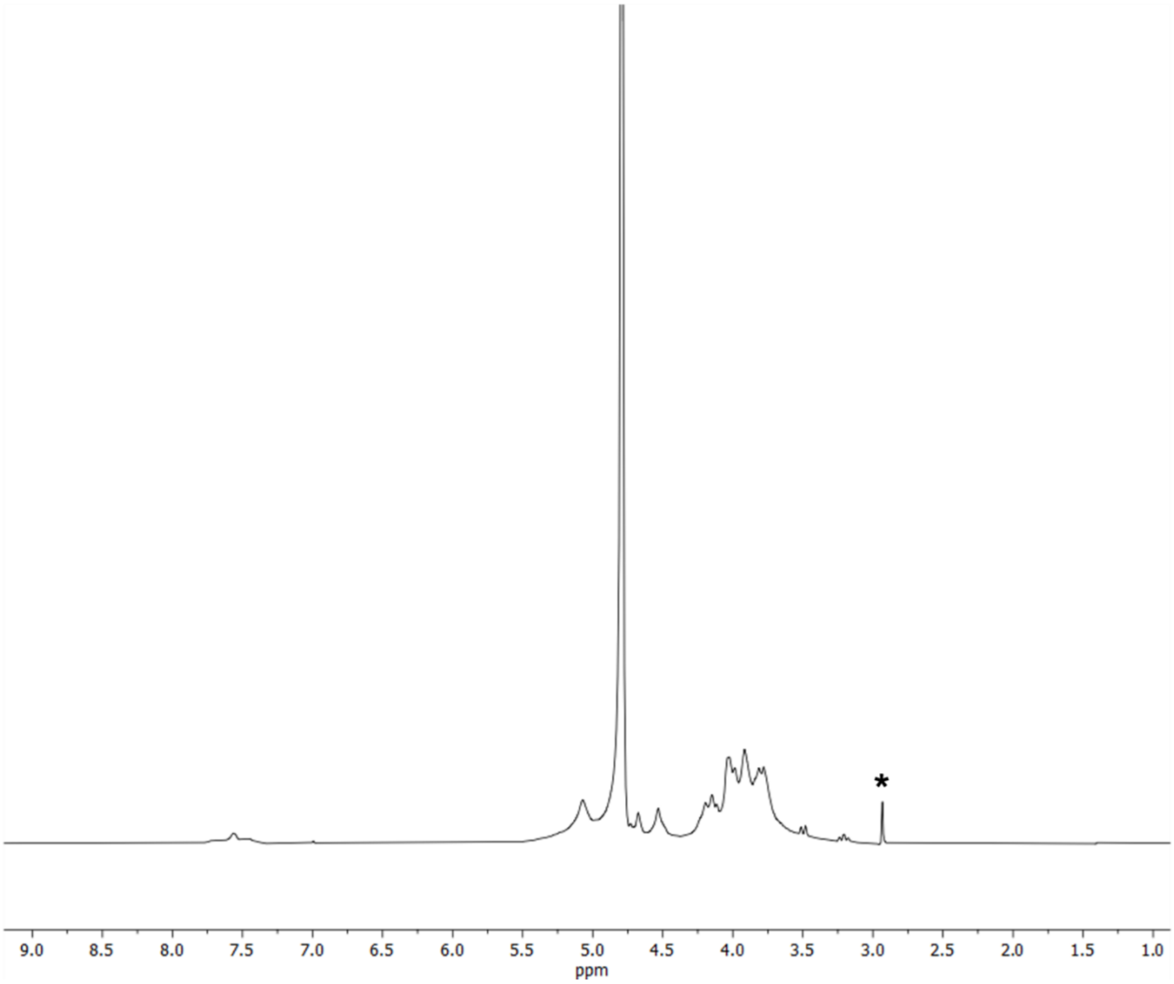


**Figure S5.** $^{1}H$ NMR spectrum (400 MHz, $D_2O$) of alginate functionalized with DBCO and FAM (Alg4-DBCO-FAM). * = DMSO.

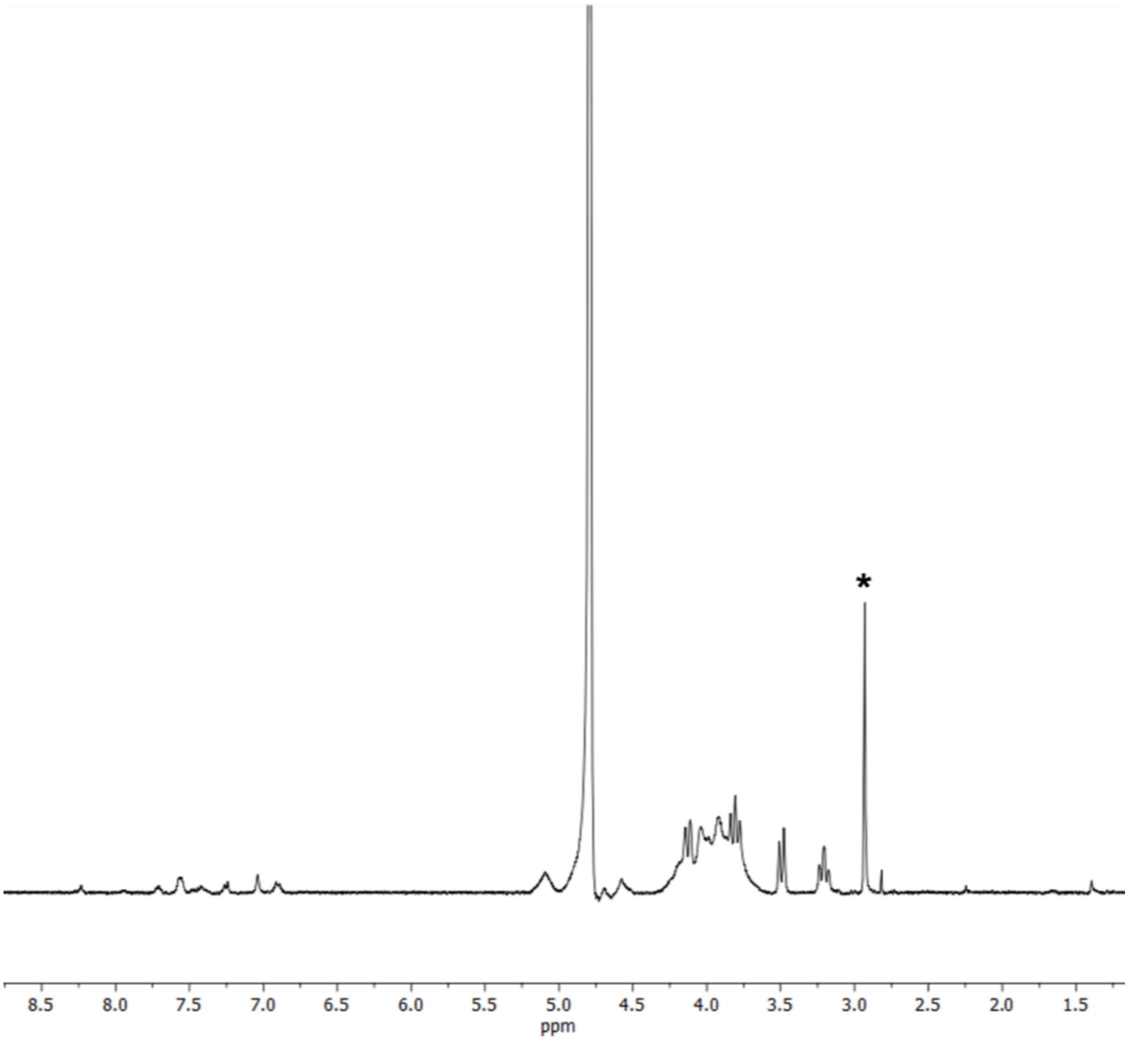


**Figure S6.** $^{1}H$ NMR spectrum (400 MHz, $D_2O$) of alginate functionalized with DBCO and FAM (Alg8-DBCO-FAM). * = DMSO.

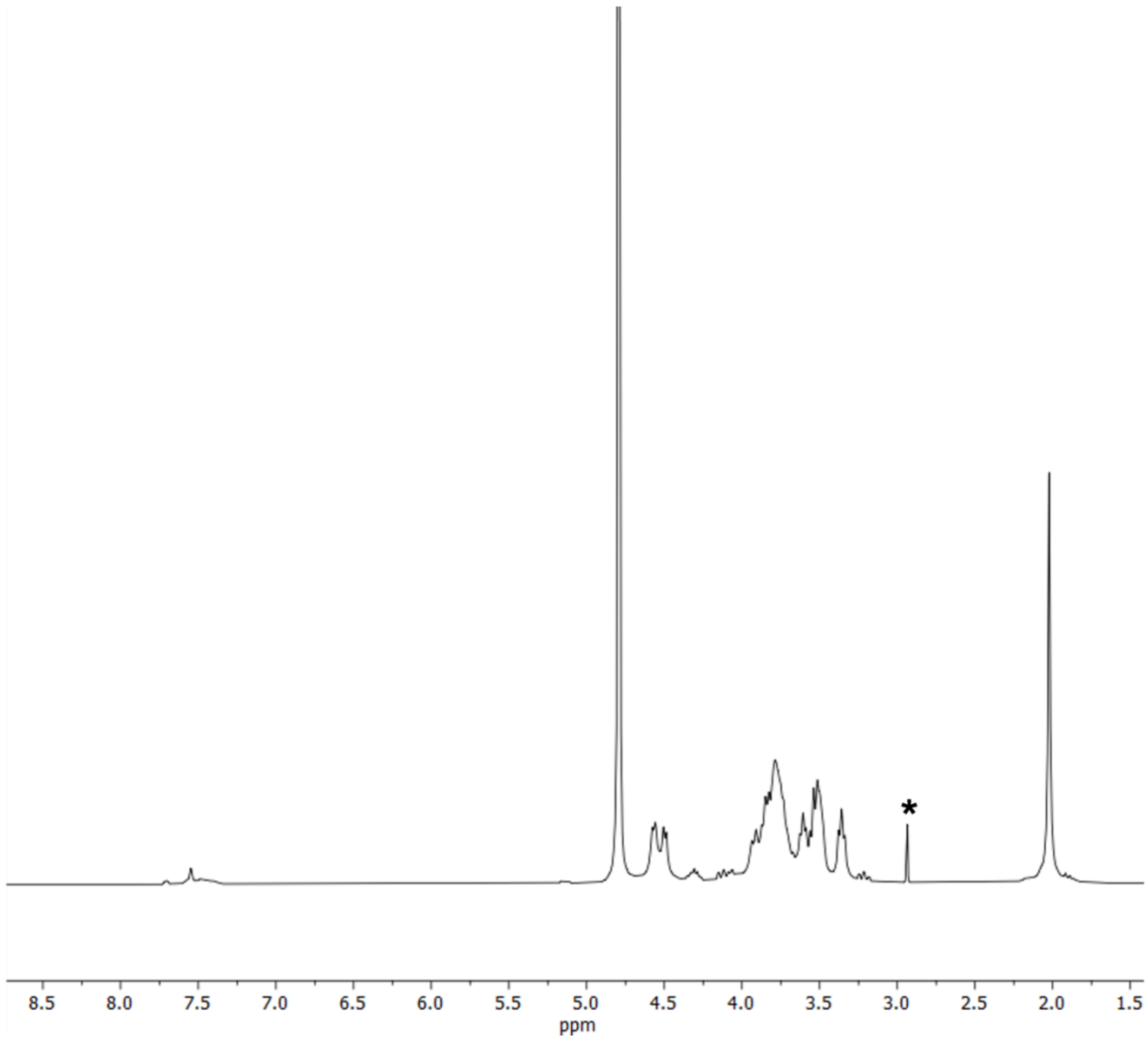


**Figure S7.** [1]H NMR spectrum (400 MHz, $D_2O$) of hyaluronic acid functionalized with DBCO and FAM (HA4-DBCO-FAM). * = DMSO.

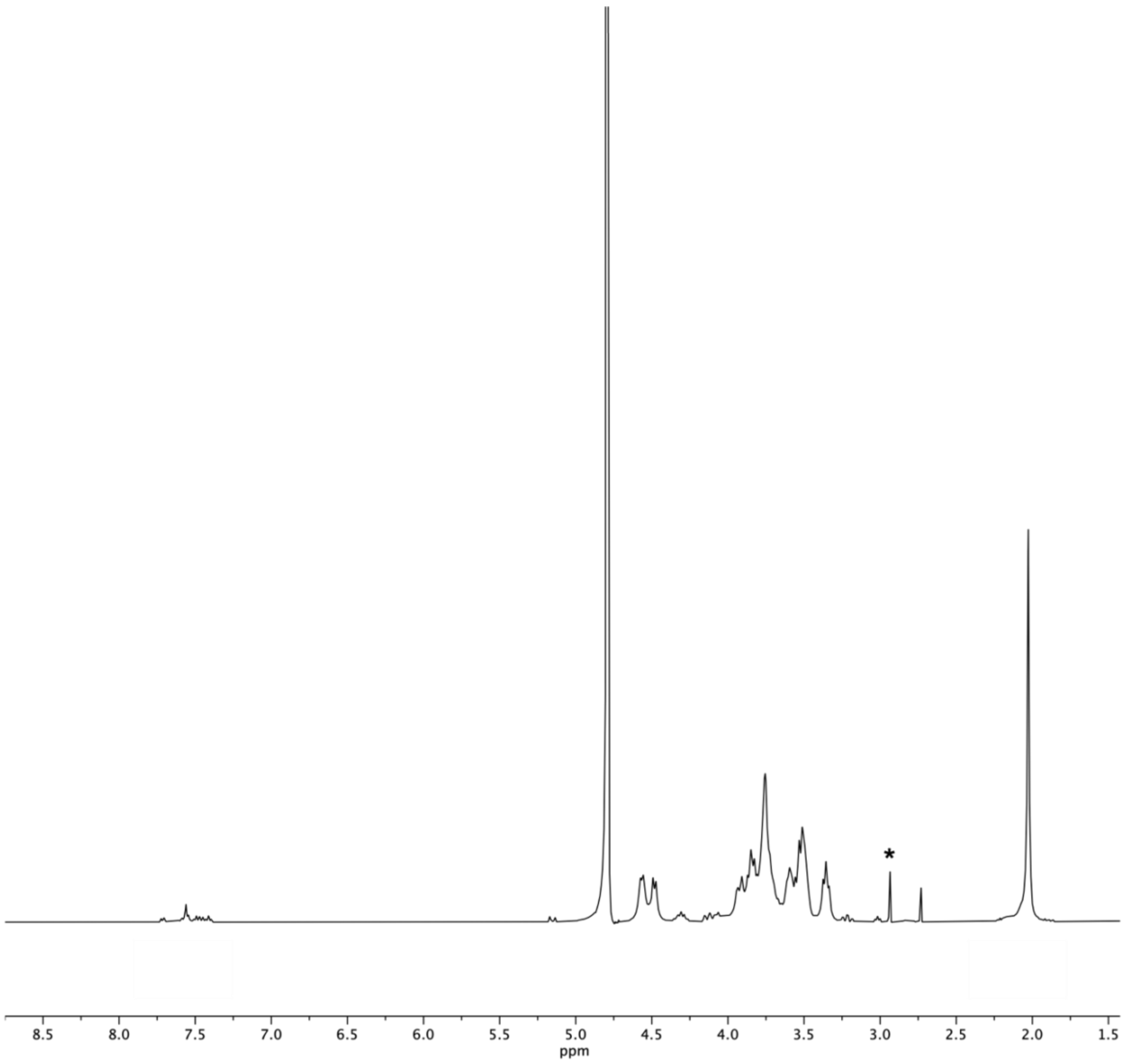


**Figure S8.** $^{1}H$ NMR spectrum (400 MHz, $D_2O$) of hyaluronic acid functionalized with DBCO and FAM (HA8-DBCO-FAM). * = DMSO.

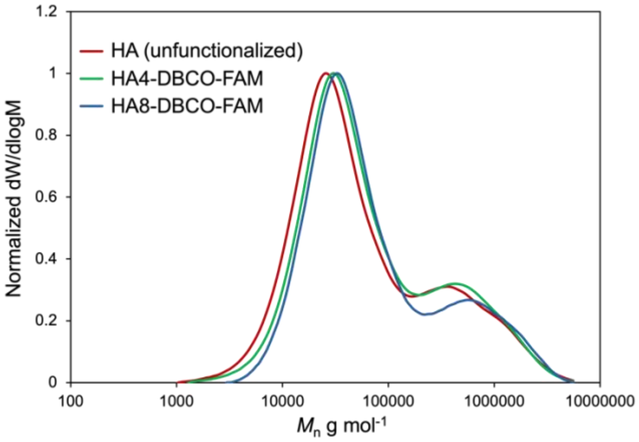


**Figure S9.** SEC of HA polymers (RI detection, elution with $H_2O$ and 20% MeOH). PEG calibration.

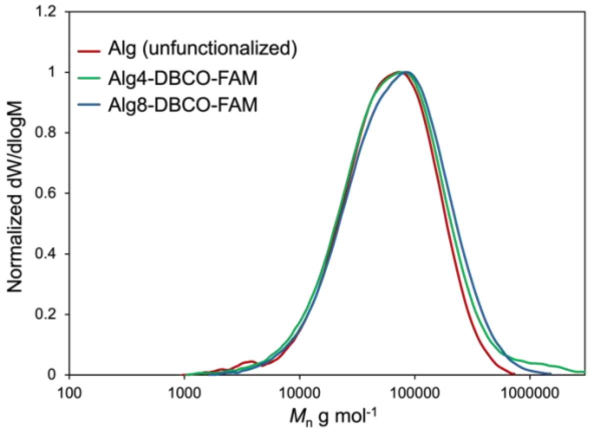


**Figure S10.** SEC of Alg polymers (RI detection, elution with $H_2O$ and 20% MeOH). PEG calibration.

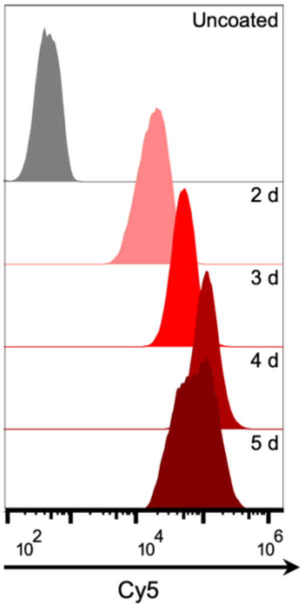


**Figure S11.** Flow cytometry graphs of HPCs treated with $Ac_4ManNAz$ for the indicated time periods (2 - 5 d) and incubated with Cy5-DBCO for 2 h.

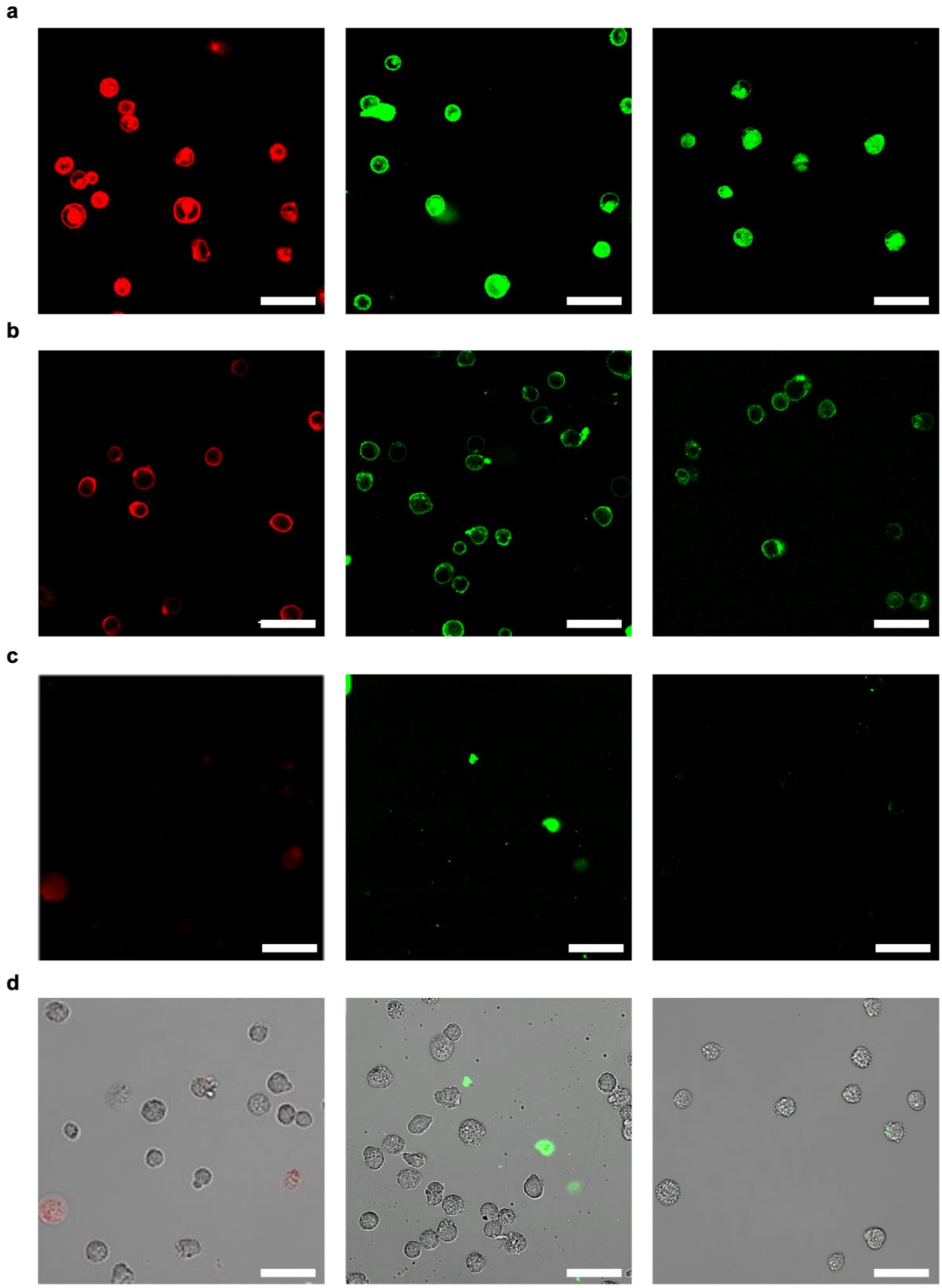


**Figure S12. a.** Representative confocal fluorescence microscopy images of azide-treated HPCs incubated at 37 °C with DBCO-Cy5, HA8 and Alg8 (left to right). **b.** Representative confocal fluorescence microscopy images of azide-treated HPCs incubated at 4 °C with DBCO-Cy5,

HA8 and Alg8 (left to right). **c.** Representative confocal fluorescence microscopy images of HPCs (not treated with azide) incubated at 4 °C with DBCO-Cy5, HA8 and Alg8 (left to right). **d.** Fluorescence and brightfield merge of HPCs (not treated with azide) incubated at 4 °C with DBCO-Cy5, HA8 and Alg8 (left to right). Scale bar = 50 µm.

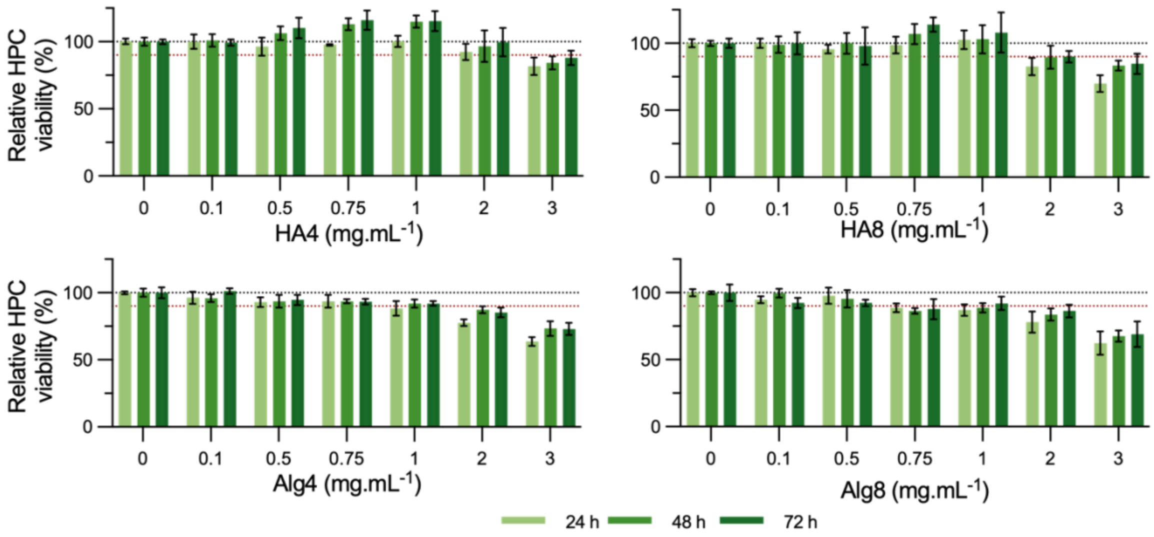


**Figure S13.** Viability of HPCs over 72 h after incubation with increasing concentrations of DBCO-functionalized polysaccharides (0 - 3 mg mL$^{-1}$) (n = 3, ± SD).

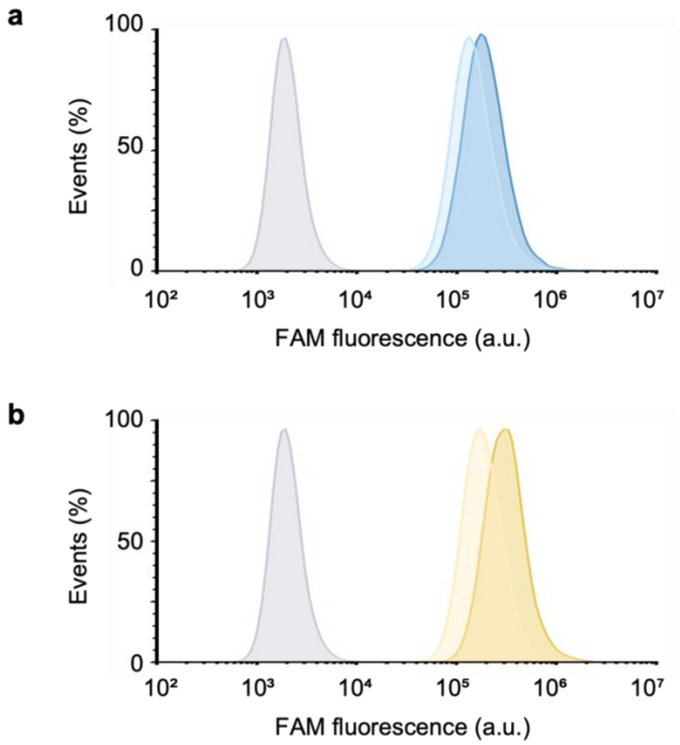


**Figure S14.** Flow cytometry analysis of HPCs uncoated (grey) and coated with HA4 and HA8 (**a**, light blue and dark blue respectively) and Alg4 and Alg8 (**b**, light yellow and dark yellow respectively).

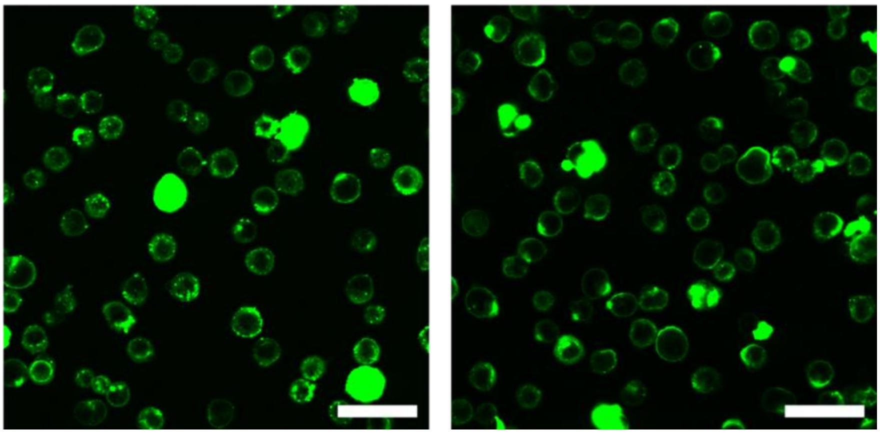

**Figure S15.** Representative confocal fluorescence microscopy images of Alg8 (left) and HA8 (right) coated cells after 2.5 h of incubation at 4 °C. Scale bar = 50 µm.

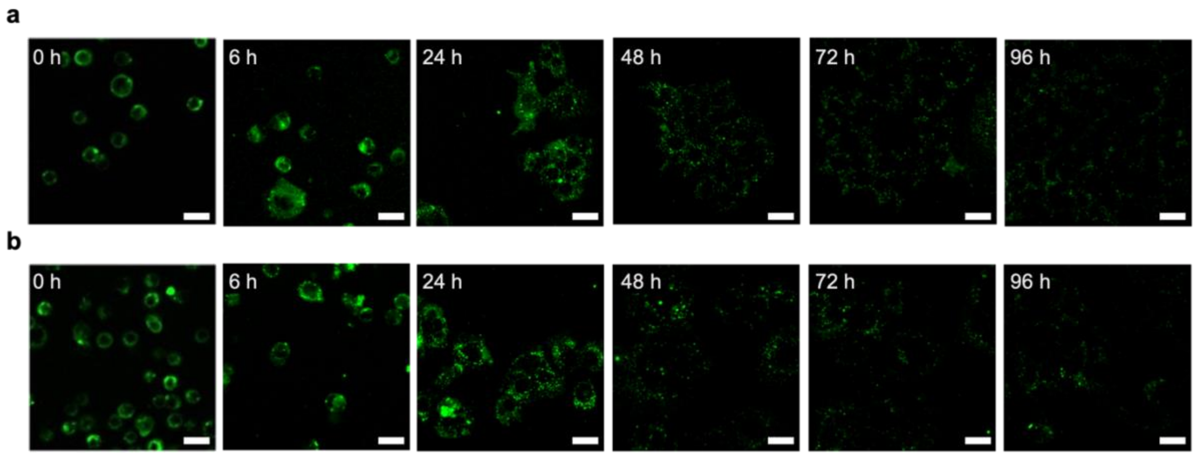


**Figure S16.** Confocal fluorescence microscopy images of HPCs in standard cell culture media after coating with HA8 **(a)** and Alg8 **(b)** captured over a period of 96 h. Scale bar = 20 µm.

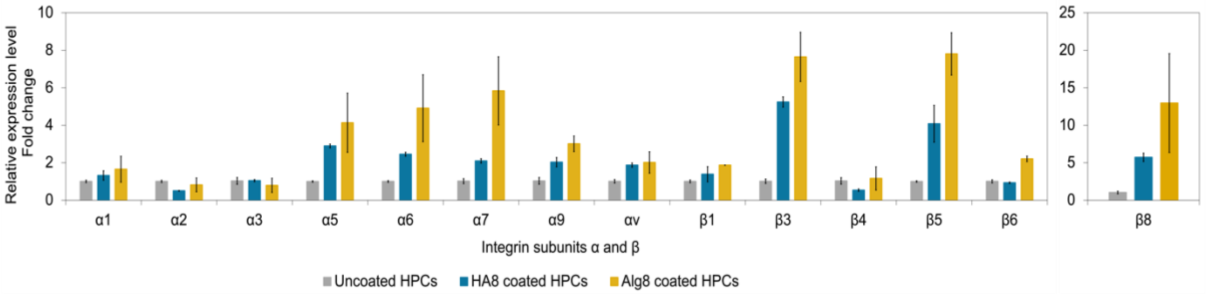


**Figure S17.** Relative expression level of different integrin genes quantified by PCR (minimum N = 2 ± standard error).

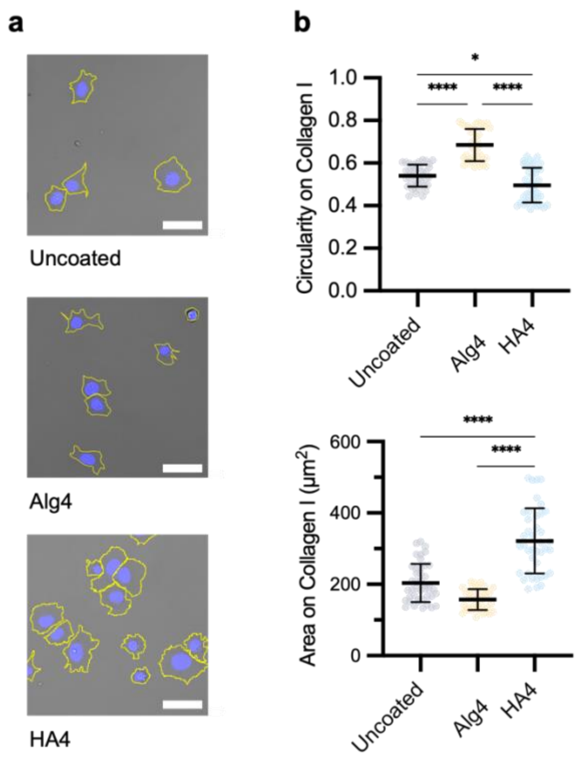


**Figure S18. a.** Representative confocal brightfield images of uncoated, Alg4, and HA4 coated HPCs left to adhere onto collagen I for 2 h. **b.** Morphological parameters for area and circularity analyzed confocal brightfield images for uncoated, Alg4, and HA4 coated HPCs after adhesion onto collagen-coated plates. Scale bar = 50 µm (N = 3 ± SD, **** P = <0.0001).

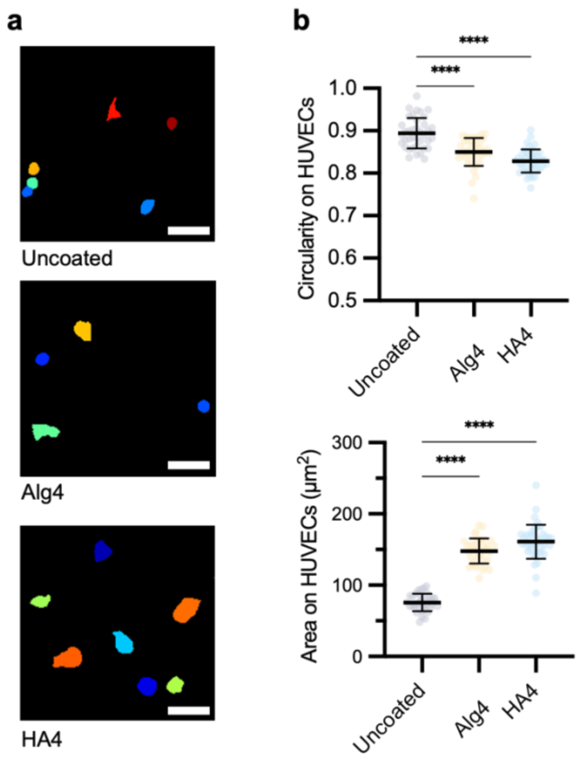


**Figure S19. a.** Representative confocal fluorescence microscopy image masks of uncoated, Alg4, and HA4 coated HPCs left to adhere onto HUVEC monolayers for 2 h. **b.** Morphological parameters for area and circularity analyzed form the image masks for uncoated, Alg4, and HA4 coated HPCs after adhesion to HUVEC monolayers. Scale bar = 50 µm (N = 3 ± SD, **** P = <0.0001).
